# Predicting the occurrence of variants in *RAG1* and *RAG2*

**DOI:** 10.1101/272609

**Authors:** Dylan Lawless, Hana Lango Allen, James Thaventhiran, NIHR BioResource-Rare Diseases Consortium, Flavia Hodel, Rashida Anwar, Jacques Fellay, Jolan E. Walter, Sinisa Savic

## Abstract

While widespread genome sequencing ushers in a new era of preventive medicine, the tools for predictive genomics are still lacking. Time and resource limitations mean that human diseases remain uncharacterised because of an inability to predict clinically relevant genetic variants. A strategy of targeting highly conserved protein regions is used commonly in functional studies. However, this benefit is lost for rare diseases where the attributable genes are mostly conserved. An immunological disorder exemplifying this challenge occurs through damaging mutations in *RAG1* and *RAG2* which presents at an early age with a distinct phenotype of life-threatening immunodeficiency or autoimmunity. Many tools exist for variant pathogenicity prediction but these cannot account for the probability of variant occurrence. Here, we present a method that predicts the likelihood of mutation for every amino acid residue in the RAG1 and RAG2 proteins. Population genetics data from approximately 146,000 individuals was used for rare variant analysis. Forty-four known pathogenic variants reported in patients and recombination activity measurements from 110 RAG1/2 mutants were used to validate calculated scores. Probabilities were compared with 98 currently known human cases of disease. A genome sequence dataset of 558 patients who have primary immunodeficiency but that are negative for RAG deficiency were also used as validation controls. We compared the difference between mutation likelihood and pathogenicity prediction. Our method builds a map of most probable mutations allowing pre-emptive functional analysis. This method may be applied to other diseases with hopes of improving preparedness for clinical diagnosis.

## Introduction

Costs associated with genomic investigations continue to reduce [1] while the richness of data generated increases. Globally, the adoption of wide scale genome sequencing implies that all new-born infants may receive screening for pathogenic genetic variants in an asymptomatic stage, pre-emptively [2]. The one dimensionality of individual genomes is now being expanded by the possibility of massive parallel sequencing for somatic variant analysis and by single-cell or lineage-specific genotyping; culminating in a genotype spectrum. In whole blood, virtually every nucleotide position may be mutated across 105 cells [3]. Mapping one’s genotype across multiple cell types and at several periods during a person’s life may soon be feasible [4]. Such genotype snapshots might allow for prediction and tracking of somatic, epigenetic, and transcriptomic profiling.

The predictive value of genomic screening highly depends on the computation tools used for data analysis and its correlation with functional assays or prior clinical experience. Interpretation of that data is especially challenging for rare human genetic disorders; candidate disease-causing variants that are predicted as pathogenic often require complex functional investigations to confirm their significance. There is a need for predictive genomic modelling with aims to provide a reliable guidance for therapeutic intervention for patients harbouring genetic defects for life-threatening disease before the illness becomes clinically significant.

The study of predictive genomics is exemplified by consideration of gene essentiality, accomplished by observing intolerance to loss-of-function variants. Several gene essentiality scoring methods are available for both the coding and non-coding genome [5]. Approximately 3,000 human genes cannot tolerate the loss of one allele [5]. The greatest hurdle in monogenic disease is the interpretation of variants of unknown significance while functional validation is a major time and cost investment for laboratories investigating rare disease.

Severe, life-threatening immune diseases are caused by genetic variations in almost 300 genes [6, 7] however, only a small percentage of disease causing variants have been characterised using functional studies. Several robust tools are in common usage for predicting variant pathogenicity. Compared to methods for pathogenicity prediction, a void remains for predicting mutation probability, essential for efficient pre-emptive validation. Our investigation aims to apply predictive genomics as a tool to identify genetic variants that are most likely to be seen in patient cohorts.

We present the first application of our novel approach of predictive genomics using Recombination activating gene 1 (RAG1) and RAG2 deficiency as a model for a rare primary immunodeficiency (PID) caused by autosomal recessive variants. *RAG1* and *RAG2* encode lymphoid-specific proteins that are essential for V(D)J recombination. This genetic recombination mechanism is essential for a robust immune response by diversification the T and B cell repertoire in the thymus and bone marrow, respectively [8, 9]. Deficiency of RAG1 [10] and RAG2 [11] in mice causes inhibition of B and T cell development. Schwarz et al. [12] formed the first publication reporting that RAG mutations in humans causes severe combined immunodeficiency (SCID), and deficiency in peripheral B and T cells. Patient studies identified a form of immune dysregulation known as Omenn syndrome [13, 14]. The patient phenotype includes multi-organ infiltration with oligoclonal, activated T cells. The first reported cases of Omenn syndrome identified infants with hypomophic RAG variants which retained partial recombination activity [15]. RAG deficiency can be measured by in vitro quantification of recombination activity [16–18]. Hypomorphic *RAG1* and *RAG2* mutations, responsible for residual V(D)J recombination activity (on average 5-30%), result in a distinct phenotype of combined immunodeficiency with granuloma and/or autoimmunity (CID-G/A) [2, 19, 20].

Human RAG deficiency has traditionally been identified at very early ages due to the rapid drop of maternally-acquired antibody in the first six months of life. A loss of adequate lymphocyte development quickly results in compromised immune responses. More recently, we have found that RAG deficiency is also found for some adults living with PID [16].

*RAG1* and *RAG2* are highly conserved genes but disease is only reported with autosomal recessive inheritance. Only 44% of amino acids in RAG1 and RAG2 are reported as mutated on GnomAD and functional validation of candidate variants is difficult [21]. Pre-emptive selection of residues for functional validation is a major challenge; a selection based on low allele frequency alone is infeasible since the majority of each gene is highly conserved. A shortened time between genetic analysis and diagnosis means that treatments may be delivered earlier. RAG deficiency may present with diverse phenotypes and treatment strategies vary. With such tools, early intervention may be prompted. Some patients could benefit from hematopoietic stem cell transplant [22] when necessary while others may be provided mechanism-based treatment [23]. Here, we provide a new method for predictive scoring that was validated against groups of functional assay values, human disease cases, and population genetics data. We present the list of variants most likely seen as future determinants of RAG deficiency, meriting functional investigation.

## Methods

### Population genetics and data sources

GnomAD (version r2.0.2) [21] was queried for the canonical transcripts of *RAG1* and *RAG2* from population genetics data of approximately 146,000 individuals; ENST00000299440 (*RAG1*) 1586 variants, GRCh37 11:36532259-36614706 and ENST00000311485 (*RAG2*) 831 variants, GRCh37 11:36597124 - 36619829. Data was filtered to contain the variant effect identifiers: frameshift, inframe deletion, inframe insertion, missense, stop lost, or stop gained. Reference transcripts were sourced from Ensembl in the FASTA format amino acid sequence for transcript RAG1-201 ENST00000299440.5 [HGNC:9831] and transcript RAG2-201 ENST00000311485.7 [HGNC:9832]. These sequences were converted to their three-letter code format using *One to Three* from the Sequence Manipulation Suite (SMS2) [24]. Combined Annotation Dependent Depletion (CADD) scores were sourced from https://cadd.gs.washington.edu/download (Nov 2018) and are reported by Kircher et al. [25]. The dataset used was "All possible SNVs" from whole genome data, from which we extracted the data for coding regions of *RAG1* and *RAG2*. We used the Human Gene Mutation Database (HGMD) from the Institute of Medical Genetics in Cardiff as a pre-defined source of known RAG deficiency cases http://www.hgmd.cf.ac.uk/ac/index.php (Feb 2019, free access version to NM_000448.2.) [26]. Data was formatted into CSV and imported into R for combined analysis with PHRED-scaled CADD scores and the main dataframe. The crystal structure render of DNA bound RAG complex was produced with data from RCSB Protein Data Bank (3jbw.pdb) [27]. Structures were visualised using the software VMD from the Theoretical and Computational Biophysics Group [28], imaged with Tachyon rendering [29], and colour mapped using our scoring method.

### Data processing

The population genetics input dataset used GnomAD variant allele frequencies and reference sequences processed as CSV files, cleaned and sorted to contain only amino acid codes, residue numbers, alternate residues, alternate allele frequencies, and a score of 0 or 1 to indicate presence or absence of variants where 1 represented none reported. An annotation column was also provided to label where multiple alternate variants existed. Statistics and calculation steps are listed in order in Supplemental Tables E3-E8.

The percentage of conserved residues was calculated (55.99% of amino acids contained no reported variants in RAG1, 55.98% in RAG2 (Table E4)). Basic protein statistics were generated using canonical reference transcript sequences of *RAG1* and *RAG2* with the SMS2 tool *Protein Stats* [24]. The resulting pattern percentage value was converted to a frequency based on the number of residues per protein to generate the residue frequency (*R*_*f*_). The *R*_*f*_ values were found for both proteins as shown in Table E5 and summarised in Table E6.

The count of variants per residue were found for both proteins and the mutation rates (*M*_*r*_) per residue were calculated as shown in Table E7. *M*_*r*_ was found by counting the number of mutations per residue in a window, sized to contain each protein individually. For genome-wide application the window size may be increased or decreased. In this case the window consisted of only the coding regions. The *M*_*r*_ values were then converted to the frequencies based on the number of residues per protein. Separate, and overlapping windows could also be used based on genome phase data and regions of linkage disequilibrium to account for non-random association of alleles at different loci; this might be particularly important for disorders with multiple genetic determinants.

The *M*_*r*_ and *R*_*f*_ multiply to give the raw mutation rate residue frequency (MRF) value (Table E8). This value is also shown in Tables 1 **and** E1. Our investigation used a Boolean score *C* to account for the presence or absence of a mutation in the general population; 0 for any variant existing in the population and 1 for conserved residues. *C* × *M*_*r*_ × *R*_*f*_, in our case, produced the MRF score for conserved residues. Figure 1 (ii) illustrates the raw MRF as a histogram and the MRF, after applying *C*, as a heatmap.

An important consideration for future application is whether to use this Boolean score or instead use a discrete variable which accounts for the true allele frequency in the general population. In the clinical setting, the likelihood of *de novo* mutations and inherited mutations have different impacts when considering recessive and dominant diseases. A patient is more likely to inherit a variant that exists even at a very low frequency than to acquire a random *de novo* mutation. Therefore, a value representing an allele frequency may be used to replace *C* in many investigations, particularity when considering variants that exist at low rates. PRHED-scaled CADD score data consisted of nucleotide level values. For comparison with MRF, the median CADD scores were averaged per codon as demonstrated in Supplemental text. A summary of data processing and analysis is illustrated in Figure E1.

### Raw data availability and analysis script

The supplemental files *“Raw_data_R_analysis_for_figures”* contains all raw data and analysis methods used to produce figures (except illustrations in Figures 1 and 6). *“data_analysis.R”* is an R script that contains the methods used to produce figures. Each of the input data CSV files are explained on first usage within the analysis script. Running *“data_analysis.R”* from within the same directory as the associated input data CSV files will replicate analysis.

### Data visualisation

For our visualisation of MRF scores, small clusters of high MRF values were of more appealing than individual highly conserved residues. Therefore, we applied a 1% average filter where values were averaged over a sliding window of N number of residues (10 in the case of RAG1, 6 in the case of RAG2). For a clear distinction of MRF clusters, a cut-off threshold was applied at the 75^*th*^ percentile (e.g. 0.0168 in RAG1) as shown in heatmaps in Figure 1 (iii) and 6. The gene heatmaps for coding regions in *RAG1* and *RAG2* (Fig. 1) were populated with (i) Boolean *C* score from population genetics data, (ii) raw MRF scores, and (iii) MRF clusters with 1% average and cut-off threshold. GraphPad Prism was used for heatmaps. The data used for heatmaps is available in Table E1 and in the supplemental R source to allow for alternative visualisations. An example of alternative output for non-R users is shown in Figure E2. Adobe Illustrator and Photoshop were used for protein domain illustrations in Figure 1 (iv).

### Validation of MRF against functional data

The recombination activity of RAG1 and RAG2 was previously measured on known or candidate pathogenic variants [16–18]. Briefly, the pathogenicity of variants in *RAG1* and *RAG2* was measured functionally *in vitro* by either expression of RAG1 and RAG2 in combination with a recombination substrate plasmid containing recombination signal sequence (RSS) sites which are targeted by RAG complex during normal V(D)J recombination, or Abelson virus-transformed Rag2−/− pro-B cells with an RSS-flanked inverted GFP cassette. Recombination events were assessed by quantitative real-time PCR using comparative CT or expression of GFP evaluated by flow cytometry, respectively. The inverse score of recombination activity (0-100%) was used to quantify pathogenicity of variants in our study. Comparison between known pathogenicity scores and MRF was done by scaling MRF scores from 0-100% (100% being highest probability of occurring as damaging). A data and analysis is summarised in Figure E1.

## Results

### RAG1 and RAG2 conservation and mutation rate residue frequency

Variant probability prediction is dependent on population genetics data. Our study queried GnomAD [21] to identify conserved residues using a Boolean score *C* of 0 (present in population) or 1 (conserved). The gene-specific mutation rate *M*_*r*_ of each residue was calculated from allele frequencies. The gene-specific residue frequency *R*_*f*_ represented the frequency of a residue occurring per gene, acquired by converting gene residue percentage (from the SMS2 tool *Protein stats*) to a frequency [24]. Together the values were used to calculate the most probable disease-causing variants which have not yet been identified in patients. We termed the resulting score a mutation rate residue frequency, where *MRF* = *C* × *M*_*r*_ × *R*_*f*_. This score represents the likelihood that a clinically relevant mutation will occur.

Figure 1 presents the most probable unidentified disease-causing variants in *RAG1/2*. Variants with a low MRF may still be damaging but resources for functional validation are best spent on gene regions with high MRF. Clusters of conserved residues are shown in Figure 1 (i) and are generally considered important for protein structure or function. However, these clusters do not predict the likelihood of mutation. Raw MRF scores are presented in Figure 1 (ii). Histograms illustrates the MRF without Boolean scoring applied and Figure 1 (iii) provides a clearer illustration of top MRF score clusters. For visualisation, a noise reduction method was applied; a sliding window was used to find the average MRF per 1% interval of each gene. The resulting scores displayed in Figure 1 (iii) contain a cut-off threshold to highlight the top scoring residues (using the 75^*th*^ percentile). Variant sites most likely to present in disease cases are identified by high MRF scoring. This model may be expanded by the addition of phenotypic or epigenetic data (**Supplemental**).

**Figure 1:**
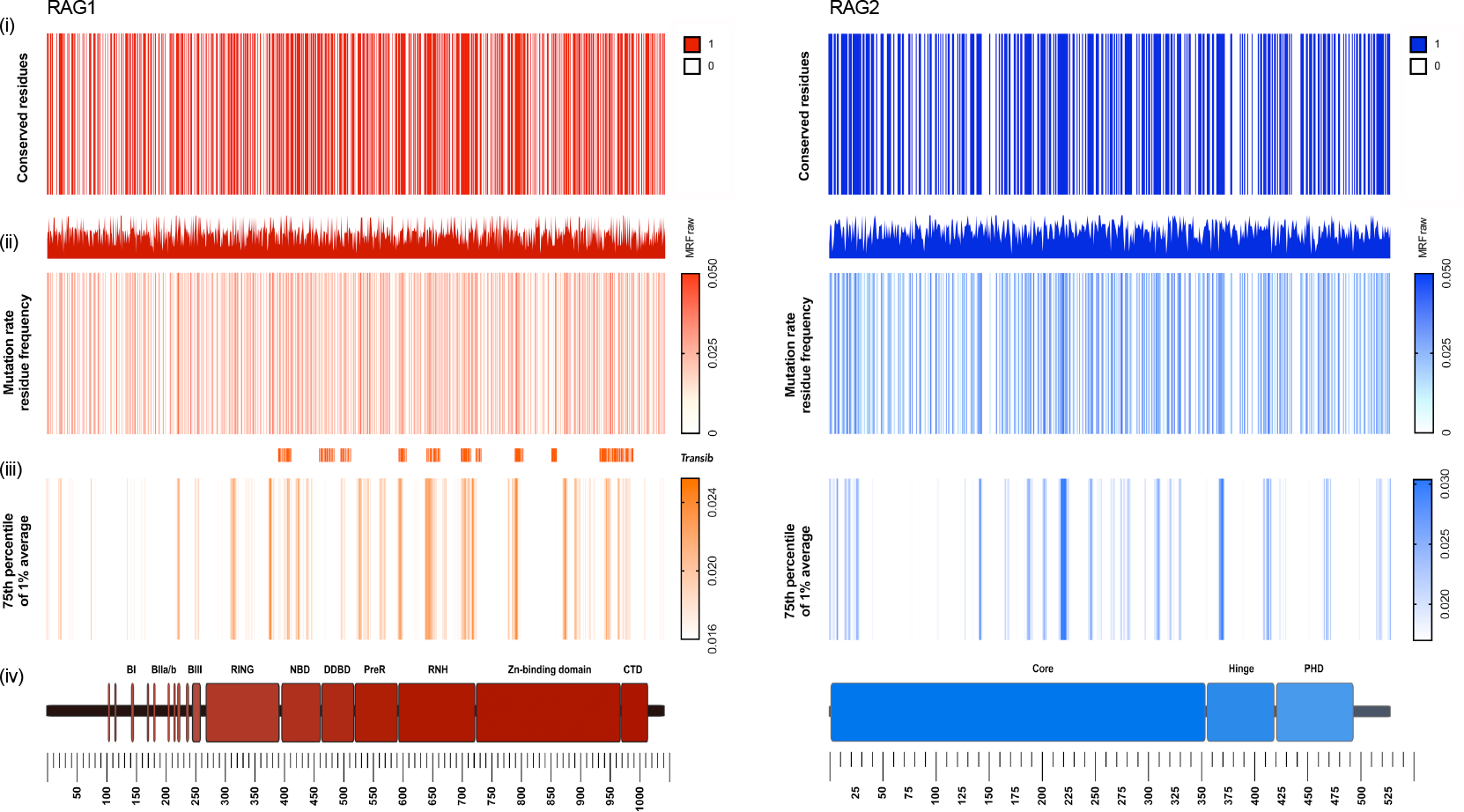
RAG1 (red, left) and RAG2 (blue, right) conservation and mutation rate residue frequency. (i) Gene conservation score; non-conserved 0 and conserved 1. Colour indicates no known mutations in humans. (ii) Histogram; raw MRF score. Heatmap; MRF prediction for conserved residues, graded 0 to 0.05 (scale of increasing mutation likelihood with human disease). (iii) Coloured bars indicate most likely clinically relevant variant clusters. MRF score averaged with 1% intervals for each gene and cut-off below 75th percentile, graded 0 to 0.03 (noise reduction method). (iv) Gene structure with functional domains. Full list of residues and scores available in Table E1.

Table E1 provides all MRF scores for both proteins. Raw data used for calculations and the list of validated residues of RAG1 and RAG2 are available in Tables E3-E8. **Summary** table 1 shows the MRF mutation likelihood score for mutations that have also been reported as tested for recombination activity in functional assays. Analysis-ready files are also available in Supplemental data along with the associated R source file to allow for alternative visualisations as shown in Figure E2.

**Table 1:**
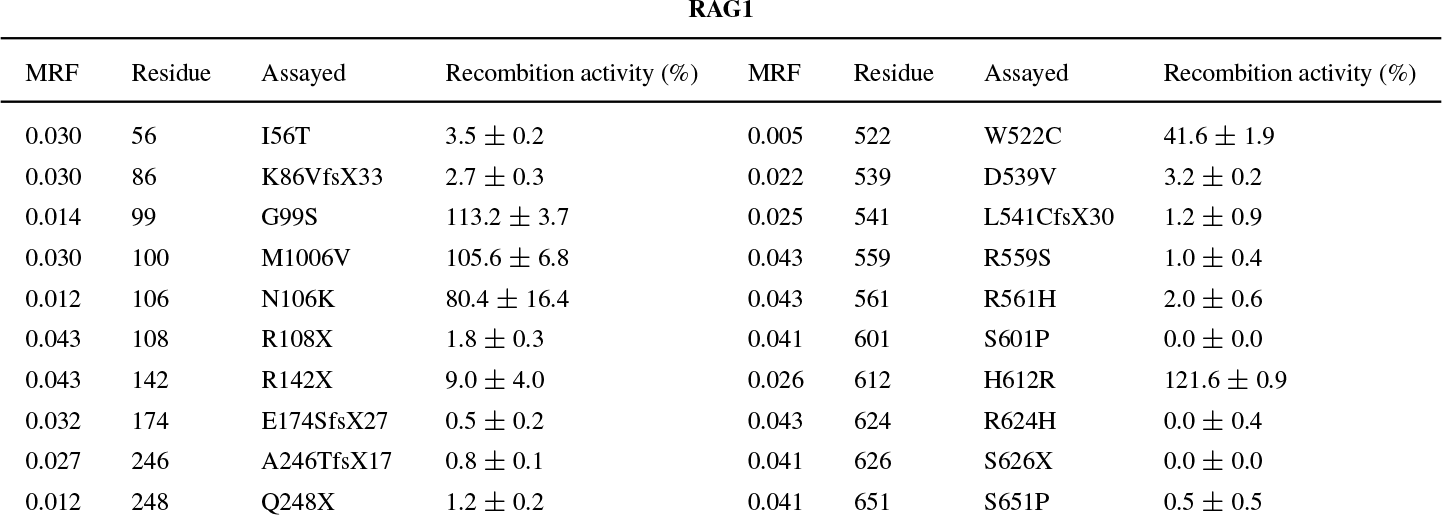
MRF likelihood scores for variants functionally assayed to date [16–18]. Increased MRF score indicate higher likelihood of occurrence. Recombination activity measured by functional assay is shown as a percentage of wildtype (% SEM). Residues with multiple mutations are shown with both alternative variants and values. *MRF*_*max*_ = 0.043 and *MRF*_*min*_ = 0.004. The full table of all protein positions can be found in Supplemental Table E1.

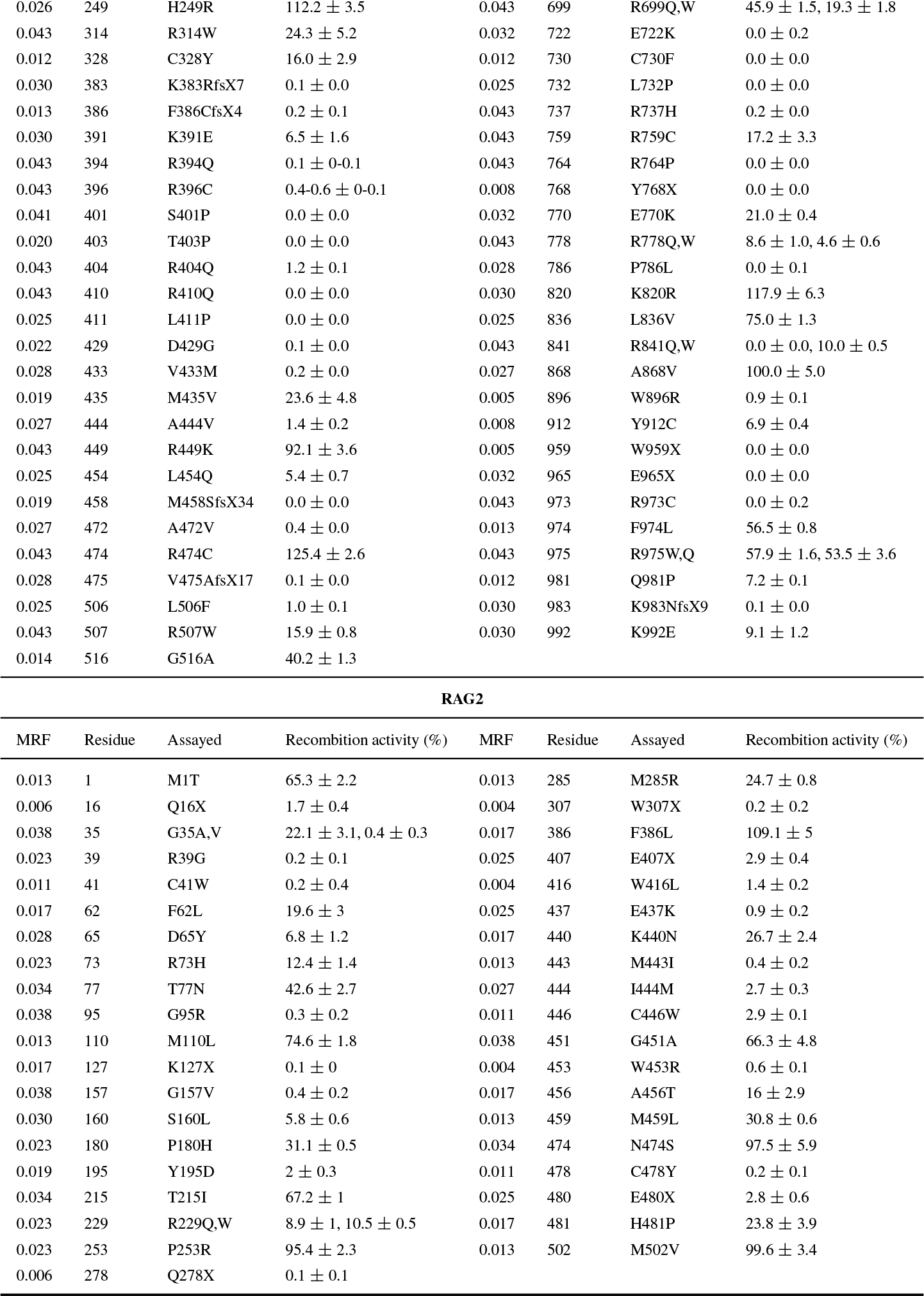

### MRF scores select for confirmed variants in human disease

We have applied MRF scores to known damaging mutations from other extensive reports in cases of human disease [12, 15, 17, 19, 20, 30–53] [*originally compiled by* Notarangelo et al. [54]]. This dataset compares a total of 44 variants. We expected that functionally damaging variants (resulting in low recombination activity in vitro) that have the highest probability of occurrence would be identified with high MRF scores. MRF prediction correctly identified clinically relevant mutations in *RAG1* and *RAG2* (Fig. 2 (i)). Variants reported on GnomAD which are clinically found to cause disease had significantly higher MRF scores than variants which have not been reported to cause disease. We observed that rare and likely mutations provided high scores while rare but unlikely or common variants had low scores (Fig. 2 (i)).

**Figure 2:**
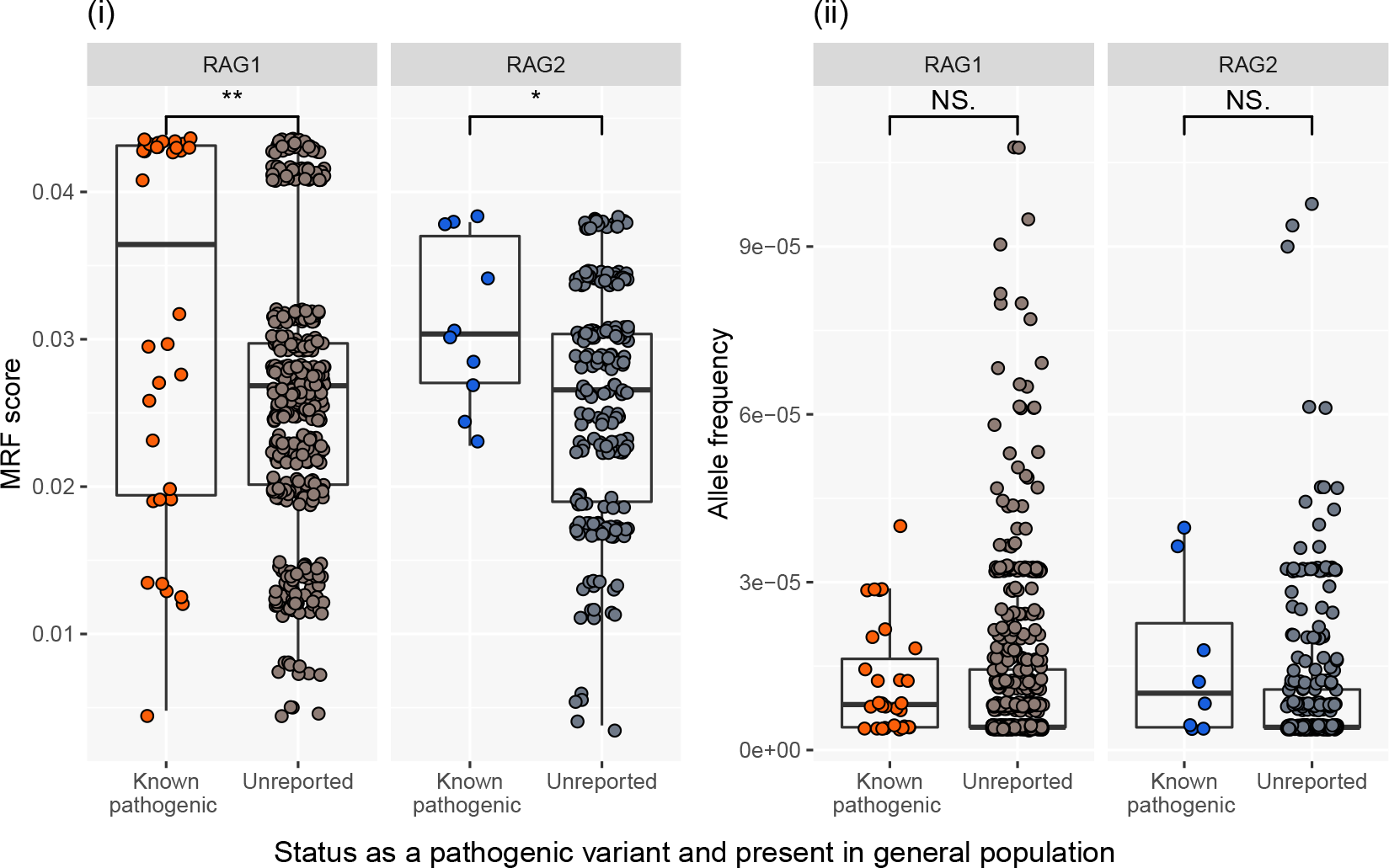
RAG1 and RAG2 MRF score predict the likelihood of mutations that are clinically relevant. (i) Known damaging variants (clinically diagnosed with genetic confirmation) reported on GnomAD have significantly higher MRF scores than unreported variants. (ii) GnomAD rare variant allele frequency <0.0001. No significant difference in allele frequency is found between known damaging and non-clinically reported variants. Unpaired t-test. RAG1 P-value 0.002** RAG2 P-value 0.0339*. MRF; mutation rate residue frequency, ns; non-significant.

Allele frequency is generally the single most important filtering method for rare disease in whole genome (and exome) sequencing experiments. Variants under pressure from purifying selection are more likely to cause disease than common variants. However, most RAG mutations are rare. Therefore, allele frequencies of rare variants reported on GnomAD cannot differentially predict the likelihood of causing disease (Fig. 2 (ii)). As such, we found no significant difference between known damaging variants and those that have not yet been reported as disease-causing. The comparison between Figure 2 (i) **and (ii)** illustrates the reasoning for the design of our method.

Many non-clinically-reported rare variants may cause disease; the MRF score identifies the top clinically relevant candidates. Based on the frequency of protein-truncating variants in the general population, *RAG1* and *RAG2* are considered to be tolerant to the loss of one allele, as indicated by their low probability of being loss-of-function intolerant (pLI) scores of 0.00 and 0.01, respectively [21]. This is particularly important for recessive diseases such as RAG deficiency where most new missense variants will be of unknown significance until functionally validated.

### Top candidate variants require validation

Functionally characterising protein activity is both costly and time consuming. RAG1 and RAG2 have now been investigated by multiple functional assays for at least 110 coding variants [16–18]. In each case, researchers selected variants in *RAG1* and *RAG2* that were potentially damaging or were identified from PID patients as the most probable genetic determinant of disease. Functional assays for RAG deficiency in those cases, and generally, measured a loss of recombination activity as a percentage of wild type function (0-100%).

Pre-emptively performing functional variant studies benefits those who will be identified with the same variants in the future, before the onset of disease complications. While more than 100 variants have been assayed in vitro, we calculated that only one-quarter of them are most probable candidates for clinical presentation. Figure 3 illustrates that while functional work targeted “hand picked” variants that were ultimately confirmed as damaging, many of them may be unlikely to arise based on population genetics data. Figure 3 presents, in increasing order, the number of potential variants based on likelihood of presentation and stacked by the number of variants per score category. Variants that have been measured for their loss of protein activity are coloured by severity. Potential variants that remain untested are coloured in grey. Only 21 of the top 66 most probable clinically relevant variants have been assayed in *RAG1*.

Supplemental Figure E3 further illustrates the individual variants which have been tested functionally (the coloured *recombination activity* subset of Fig 3). We compared predicted MRF scores to assay measurements for 71 *RAG1* and 39 *RAG2* mutants. Most mutations tested showed severe loss of protein function (bottom panel of Supplemental Figure E3), while the likelihood each mutation occurring in humans varied significantly (top panels).

If MRF scoring was used in the same cases pre-emptively, the loss of investment would be minimal; only 8 variants out of 71 mutants tested had an above-average MRF score while being measured as functionally benign (a rate of 11.27%). RAG2 had only 3 out of 39 variants (7.69%) with an above-average MRF score while functionally benign. For the expended resources, approximately 30% more top candidates would have been tested in place of unlikely and functionally non-damaging mutations. However, the true measurement of accuracy is limited in that very few of the most likely clinically relevant variants predicted by MRF scoring have been tested to date.

**Figure 3:**
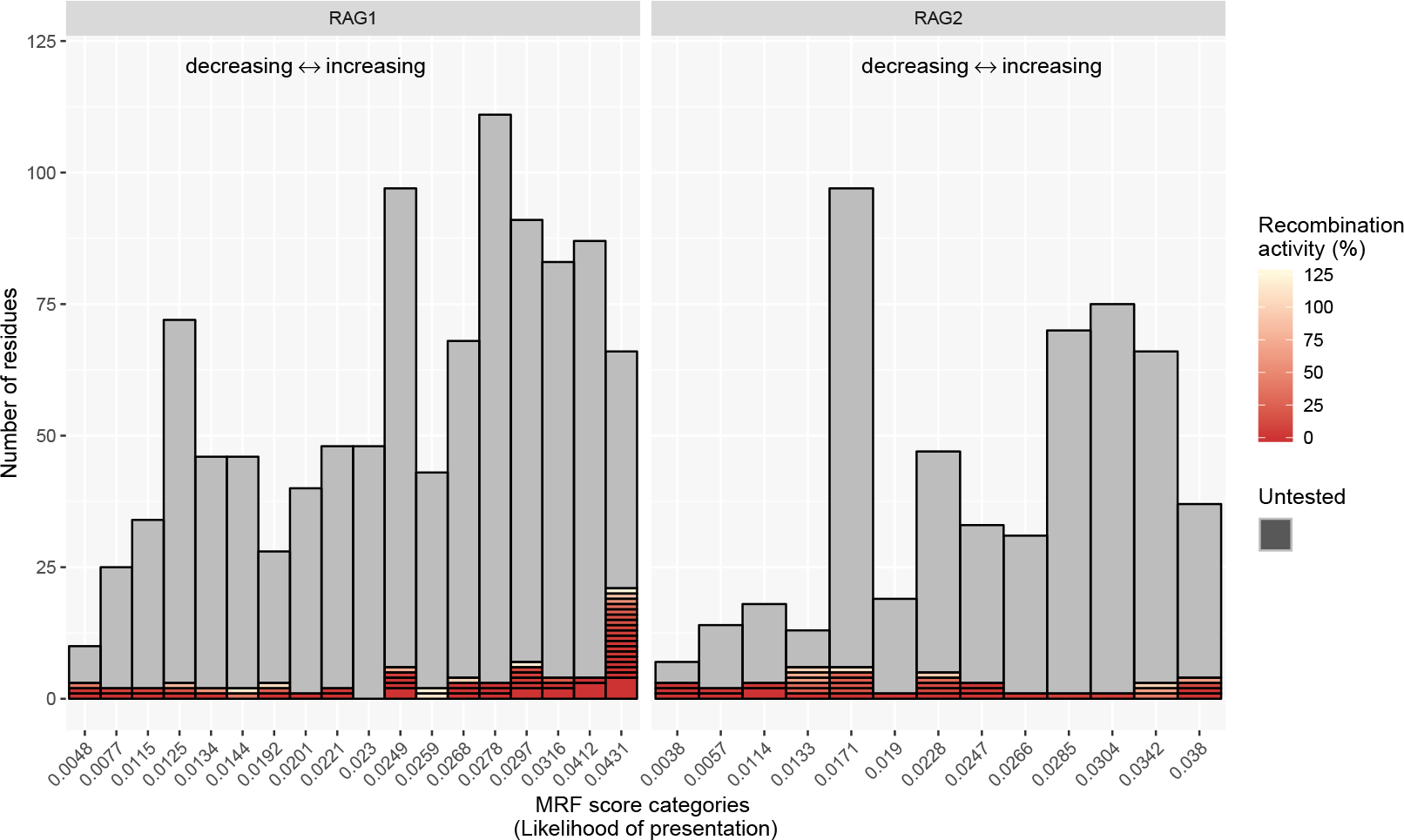
RAG1 and RAG2 MRF score categories and variants assayed to date. Protein residues are ranked and stacked into categories based on their MRF score. High scores (0.043 and 0.038 in RAG1 and RAG2, respectively) represent a greater mutation likelihood. Functional assays have measured recombination activity (as its inverse; % loss of activity) in a total of 110 mutants. The severity of protein loss of function is represented by a red gradient. Residues that have not been functionally tested are shown in grey. While many protein residues are critical to protein function, their mutation is less probable than many of the top MRF candidates. Data further expanded in Figure E3. MRF; mutation rate residue frequency.

### False positives in Transib domains do not negatively impact prediction

Adaptive immunity is considered to have evolved through jawed vertebrates after integration of the RAG transposon into an ancestral antigen receptor gene [55, 56]. The *Transib* transposon is a 600 amino acid core region of RAG1 that targets RSS-like sequences in many invertebrates. A linked *RAG1/RAG2* was shown in the lower dueterosome (sea urchin), indicating an earlier common ancestor than the invertebrate [57], and more recently, a recombinatorially active RAG transposon (ProtoRAG) was found in the lower chordate amphioxus (or lancelet); the most basal extant chordate and a “living fossil of RAG” [58].

**Figure 4:**
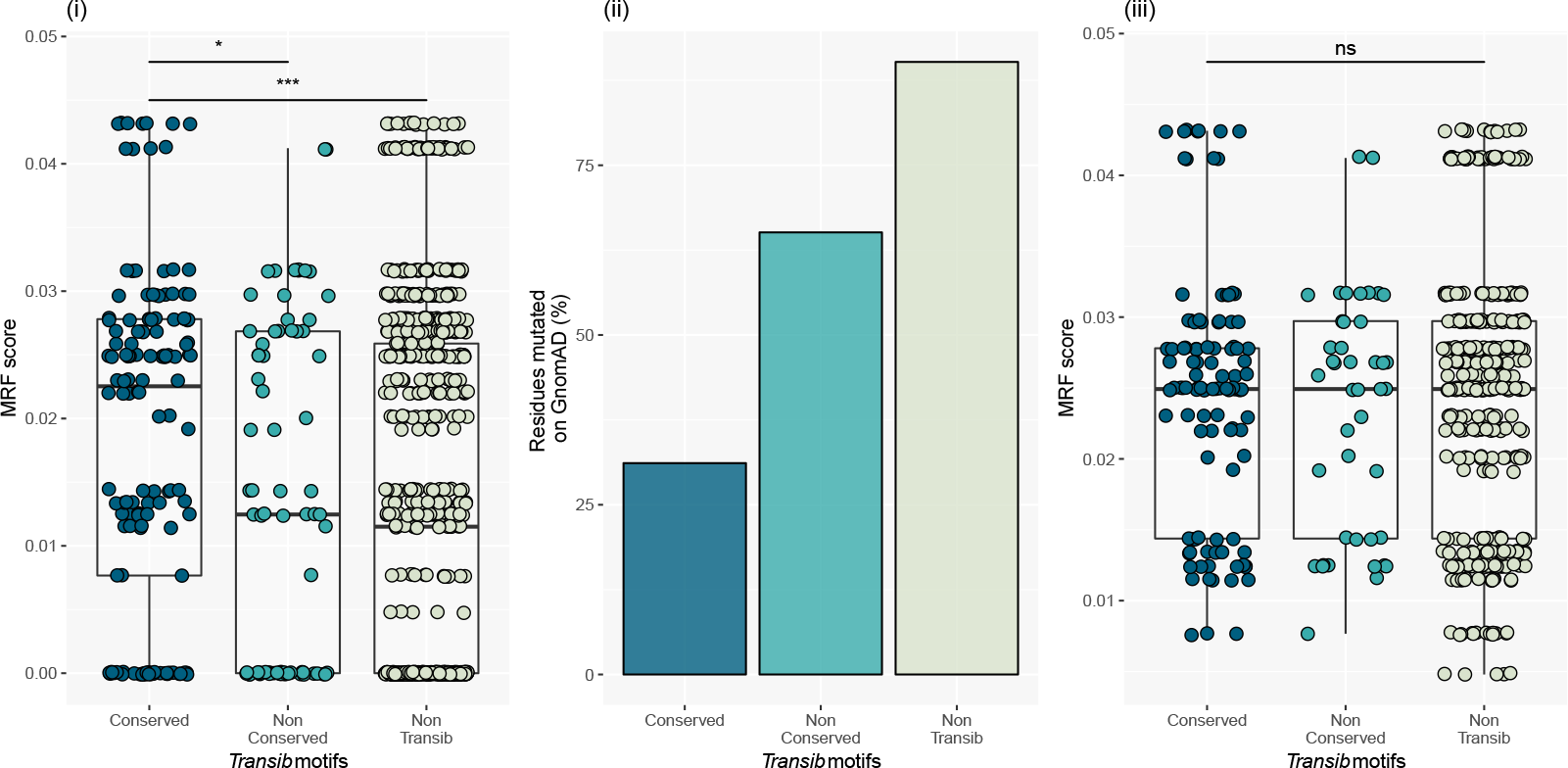
False positives in *Transib* domains do not worsen probability prediction. The *Transib* domains contain critical conserved protein residues. (i) False positives were simulated by scoring *Transib* domain MRF without omitting Boolean conservation weight *C* = 0. (ii) Allele frequencies on GnomAD had conservation levels inversely proportional to simulated false-positive MRF scoring. (iii) When testing for all Boolean component *C* > 0 after MRF calculation the effect of false positives remained non-significant, illustrating the non-negative impact of MRF for prediciting the mutation. Unpaired t-test, * P = 0.0195, *** P < 0.0001. MRF; mutation rate residue frequency, ns; non-significant.

A set of conserved motifs in core *RAG1* are shared with the *Transib* transposase, including the critical DDE residue catalytic triad (residues 603, 711, and 965) [59]. Ten *RAG1* core motifs are conserved amongst a set of diverse species including human [59]. This evolutionarily conserved region is considered as most important to protein function. Therefore, we chose this region to determine if MRF scoring would have a negative impact if mutations were falsely predicted as clinically important. To assess the influence of a false positive effect on prediction, the MRF scores for conserved residues in this group were compared to GnomAD allele frequencies. Figure 4 (i) plots the MRF (without omitting the Boolean component *C* = 0) for conserved *Transib* motif residues, non-conserved *Transib* motif residues, and non-*Transib* residues. Figure 4 (ii) shows the percentage of these which were reported as mutated on GnomAD. By accounting for unreported variants by applying *C* > 0, the resulting effect on incorrectly scoring MRF in the conserved *Transib* motifs remained neutral.

### MRF predicts RAG deficiency amongst PID patients harbouring rare variants

We have previously measured the recombination activity of RAG1 and RAG2 disease-causing variants in several patients [16]. We have compiled our own and other functional assay data from Lee et al. [17] and Tirosh et al. [18] to produce a panel of recombination activity measurements for coding variants in both *RAG1* and *RAG2*. RAG deficiency was measured as the level of recombination potential produced by the protein complex. Each method of investigation simulated the efficiency of wild-type or mutant proteins expressed by patients for their ability to produce a diverse repertoire of T-cell receptor (TCR) and B-cell receptor (BCR) and coding for immunoglobulins. In functional experiments, mutant proteins were assayed for their ability to perform recombination on a substrate which mimics the RSS of TCR and BCR in comparison to wild-type protein complex (as % SEM).

By gathering confirmed RAG deficiency cases, we compiled the MRF scores for 43 damaging *RAG1* variants in 77 PID cases and 14 damaging *RAG2* variants in 21 PID cases (MRF scores spanning over 22 categories). To test our method against a strong control group, we identified coding variants in patients with PID where RAG deficiency due to coding variants has been ruled out as the cause of disease. We obtained *RAG1/2* variants in 558 PID patients who had their genomes sequenced as part of the NIHR BioResource - Rare Diseases study [16]. Filtering initially identified 32 variants in 166 people. This set was trimmed to contain only rare variants; 29 variants over 26 MRF scoring categories from 72 cases of non-RAG-deficient PID. Linear regression on this control group produced negative or near-zero slopes for *RAG1* and *RAG2*, respectively. The same analysis for known-damaging mutations in disease cases had a significant prediction accuracy for *RAG1*. Analysis for *RAG2* was not significant. However, the sample size to date may be too small to significantly measure *RAG2* MRF scoring although a positive correlation was inferred in Figure 5 [60]. R source and raw data can be found in supplemental material.

**Figure 5:**
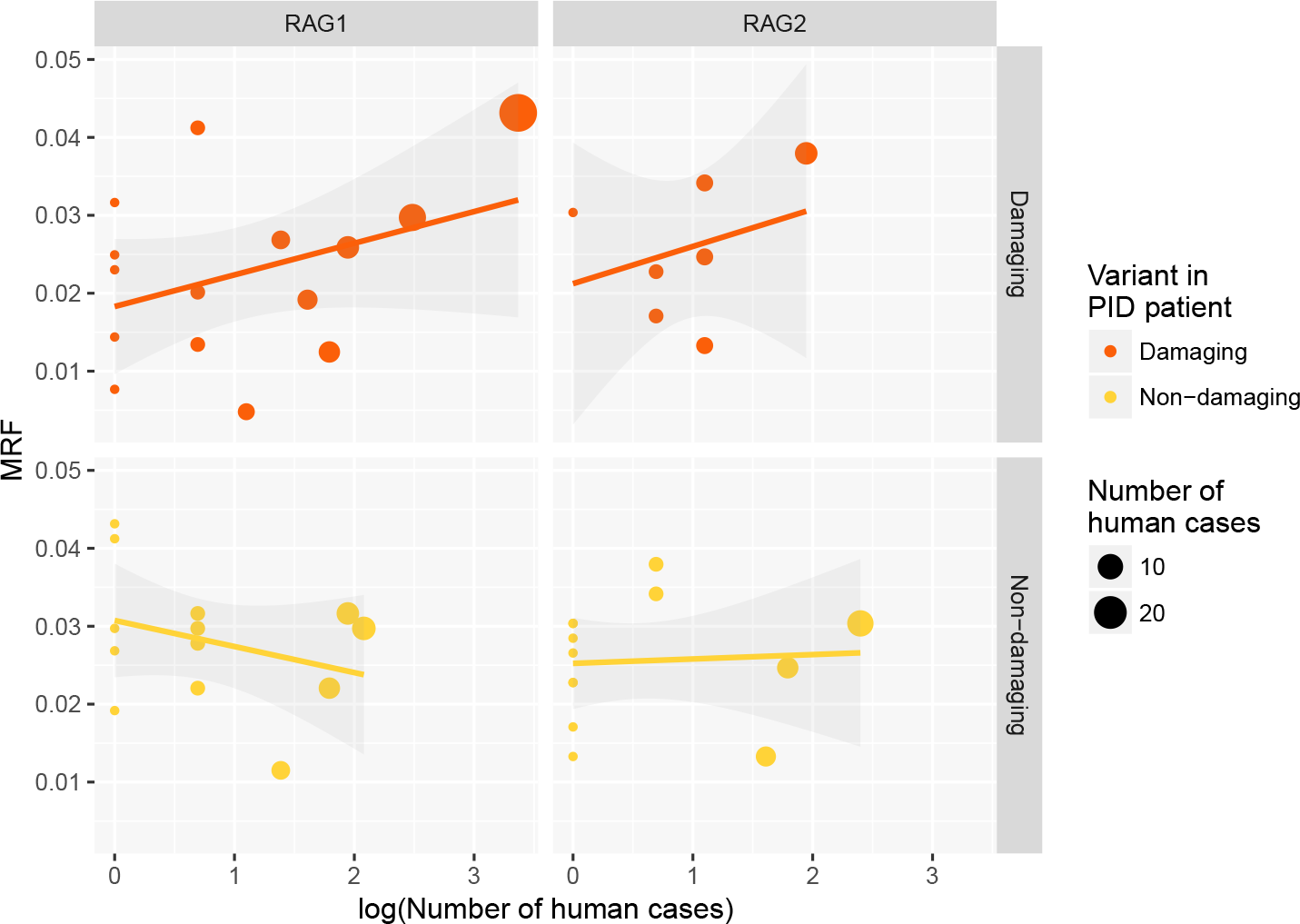
A linear regression model of RAG1/2 MRF scoring in cases of primary immune deficiency. MRF prediction correlates with clinical presentation. Damaging variants identified in confirmed RAG deficiency cases. Non-damaging variants sourced from cases of PID with rare variants but not responsible for disease. (Slopes of RAG1: Damaging: 0.0008* (± 0.0004) P<0.05, intercept 5.82e-05 ***, Non-damaging: −0.0007 (± 0.001). Slopes of RAG2; Damaging: 0.0023 (± 0.0018), intercept 0.0312 *, Non-damaging 0.0001 (± 0.0008). Source data and script in supplemental material).

### MRF supplements pathogenicity prediction tools for translational research

CADD scoring [25] is an important bioinformatics tool that exemplifies pathogenicity prediction. While CADD is a valuable scoring method, its purpose is not to predict likelihood of variation. Similarly, MRF scoring is not a measure of pathogenicity. MRF scoring may be complemented by tools for scoring variant deleteriousness. We compare MRF to the PHRED-scaled CADD scores for all possible SNV positions in *RAG1* (Fig. 6) illustrating that pathogenicity prediction cannot account for mutation probability. Combining both methods allows researchers to identify highly probable mutations before querying predicted pathogenicity.

**Figure 6:**
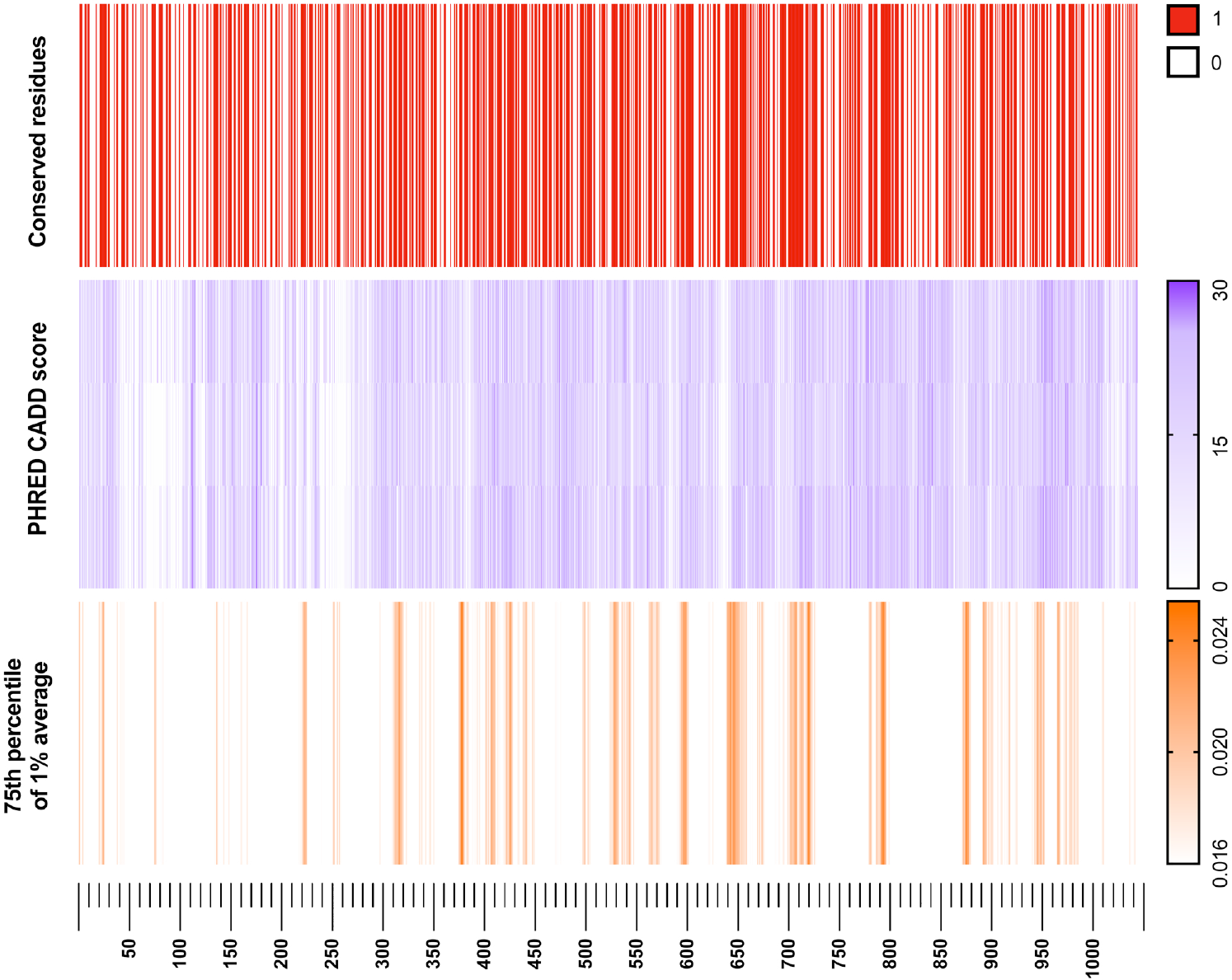
*RAG1* PHRED-scaled CADD score versus GnomAD conservation rate and MRF score. Allele frequency conservation rate (**top**) is vastly important for identifying critical structural and functional protein regions. The impact of mutation in one of these conserved regions is often estimated using CADD scoring (**middle**). CADD score heatmap is aligned by codon and separated into three layers for individual nucleotide positions. The MRF score (**bottom**)(visualised using the 75^*th*^ percentile with 1% averaging) highlights protein regions which are most likely to present clinically and may require pre-emptive functional investigation.

To further develop this concept, we first annotated variants with MRF likelihood scores and pathogenic prediction PHRED-scaled CADD scores (Figure 7), and secondly, performed a manual investigation of the clinical relevance of top candidates (Table E2). We used HGMD as an unbiased source of known RAG deficiency cases in both instances. CADD score was very successful at predicting the pathogenicity of a variant, (a high-density cluster of variants with CADD scores >25) as shown in **red** in Figure 7 (i). At about the same rate, CADD score also predicted variants as pathogenic that are, to date, unreported (as **pink** in Fig. 7 (i)). Indeed, those unreported variants may very well be pathogenic. However, the likelihood of each mutation varies. As such, we developed the MRF score to account for that likelihood. As expected, the likelihood of mutations occurring that were unreported was low according to MRF (Fig. 7 (ii), **pink**), while the mutations which did occur were highly enriched in at high MRF scores (Fig. 7 (ii), **red** high-density cluster >0.043). Combining mutation prediction (MRF) with pathogenicity prediction (tools like CADD) increases the accuracy of pre-emptively targeting clinically relevant variants. Figure 7 (iii) shows that while the number of variants presented to date is relatively small, they already account for 36% of the top MRF score candidates.

**Figure 7:**
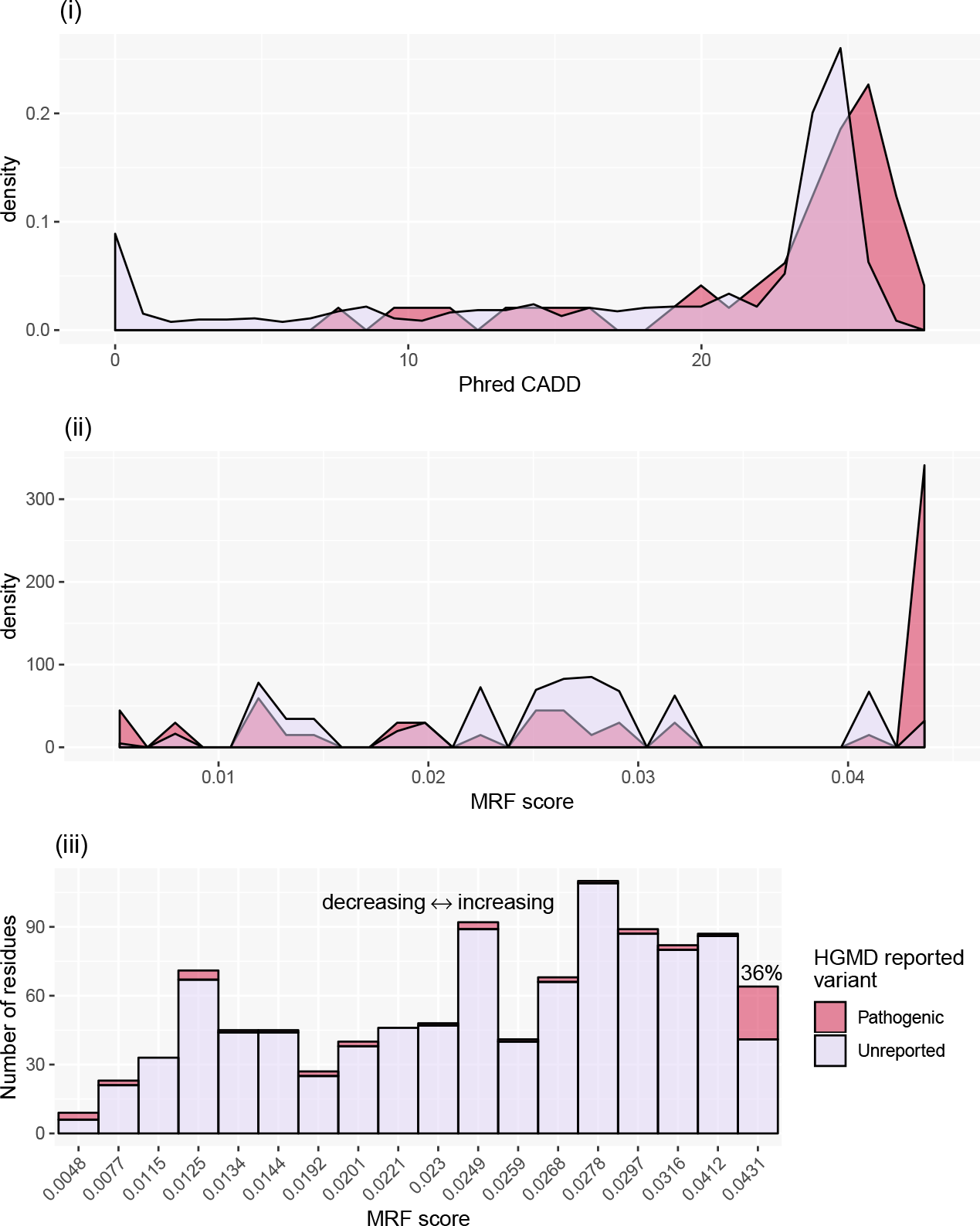
*RAG1* PHRED-scaled CADD score versus MRF score against HGMD data. (i) A high CADD score is a predictor of deleteriousness. Both reported (red) and non-reported residues (pink) have a high density of high CADD score. (ii) MRF scores only show a high-density cluster for high-likelihood variants, reflected by the high MRF score observed for known RAG deficiency variants. The number of pathogenic variants is outweighed by conserved residues; (i-ii) shows density of scores to normalise between groups. AUC overlap difference in CADD score of 21.43% and MRF score of 74.28% (above intersects >22.84 and >0.0409, in *(i-ii)* respectively). (iii) The number of residues per MRF category shows that disease reported on HGMD accounts for 36% of top MRF candidates. AUC; area under curve, CADD; Combined Annotation Dependent Depletion, HGMD; Human Gene Mutation Database.

## Discussion

Determining disease-causing variants for functional analysis typically aims to target conserved gene regions. On GnomAD 56% of *RAG1* (approx. 246,000 alleles) is conserved with no reported variants. Functional validation of unknown variants in genes with this level purifying selection is generally infeasible. Furthermore, we saw that a vast number of candidates are “predicted pathogenic” by commonly used pathogenicity tools, which may indeed be damaging but unlikely to occur. To overcome the challenge of manual selection we quantified the likelihood of mutation for each candidate variant.

Targeting clearly defined regions with high MRF scores allows for functional validation studies tailored to the most clinically relevant protein regions. An example of high MRF score clustering occurred in the RAG1 catalytic RNase H (RNH) domain at p.Ser638-Leu658 which is also considered a conserved *Transib* motif.

While many hypothetical variants with low MRF scores may be uncovered as functionally damaging, our findings suggest that human genomic studies will benefit by first targeting variants with the highest probability of occurrence (gene regions with high MRF). Table E1 lists the values for calculated MRFs for RAG1 and RAG2.

We have presented a basic application of MRF scoring for RAG deficiency. The method can be applied genome-wide. This can include phenotypically derived weights to target candidate genes or tissue-specific epigenetic features. In the state presented here, MRF scores are used for pre-clinical studies. A more advanced development may allow for use in single cases. During clinical investigations using personalised analysis of patient data, further scoring methods may be applied based on disease features. A patient phenotype can contribute a weight based on known genotype correlations separating primary immunodeficiencies or autoinflammatory diseases [6]. For example, a patient with autoinflammatory features may require a selection that favors genes associated with proinflammatory disease such as *MEFV* and *TNFAIP3*, whereas a patient with mainly immunodeficiency may have preferential scoring for genes such as *BTK* and *DOCK8*. In this way, a check-list of most likely candidates can be confirmed or excluded by whole genome or panel sequencing. However, validation of these expanded implementations requires a deeper consolidation of functional studies than is currently available.

Havrilla et al. [61] have recently developed a method with similar possible applications for human health mapping constrained coding regions. Their study employed a method that included weighting by sequencing depth. Similarly, genome-wide scoring may benefit from mutation significance cut-off, which is applied for tools such as CADD, PolyPhen-2, and SIFT [62]. We have not included an adjustment method as our analysis was gene-specific but implementation is advised when calculating genome-wide MRF scores.

The MRF score was developed to identify the top most probable variants that have the potential to cause disease. It is not a predictor of pathogenicity. However, MRF may contribute to disease prediction; a clinician may ask for the likelihood of RAG deficiency (or any other Mendelian disease of interest) prior to examination (**Supplemental**).

Predicting the likelihood of discovering novel mutations has implications in genome-wide association studies (GWAS). Variants with low minor allele frequencies have a low discovery rate and low probability of disease association [63], an important consideration for rare diseases such as RAG deficiency. An analysis of the NHGRI-EBI catalogue data highlighted diseases whose average risk allele frequency was low [63]. Autoimmune diseases had risk allele frequencies considered low at approximately 0.4. Without a method to rank most probable novel disease-causing variants, it is unlikely that GWAS will identify very rare disease alleles (with frequencies <0.001). It is conceivable that a number of rare immune diseases are attributable to polygenic rare variants. However, evidence for low-frequency polygenic compounding mutations will not be available until large, accessible genetics databases are available, exemplified by the NIHR BioResource Rare Diseases study [16]. An interesting consideration when predicting probabilities of variant frequency, is that of protective mutations. Disease risk variants are quelled at low frequency by negative selection, while protective variants may drift at higher allele frequencies [64].

The cost-effectiveness of genomic diagnostic tests is already outperforming traditional, targeted sequencing [1]. Even with substantial increases in data sharing capabilities and adoption of clinical genomics, rare diseases due to variants of unknown significance and low allele frequencies will remain non-actionable until reliable predictive genomics practices are developed. Bioinformatics as a whole has made staggering advances in the field of genetics [65]. Challenges that remain unsolved, hindering the benefit of national or global genomics databases, include DNA data storage and random access retrieval [66], data privacy management [67], and predictive genomics analysis methods. Variant filtration in rare disease is based on reference allele frequency, yet the result is not clinically actionable in many cases. Development of predictive genomics tools may provide a critical role for single patient studies and timely diagnosis [23].

## Conclusion

We provide a list of amino acid residues for RAG1 and RAG2 that have not been reported to date, but are most likely to present clinically as RAG deficiency. This method may be applied to other diseases with hopes of improving preparedness for clinical diagnosis.

## Acknowledgements

We gratefully acknowledge the participation of all NIHR BioResource volunteers, and thank the NIHR BioResource centres and staff for their contribution. We thank the National Institute for Health Research and NHS Blood and Transplant. The views expressed are those of the author(s) and not necessarily those of the NHS, the NIHR or the Department of Health.

## Ethics statement

The study was performed in accordance with the Declaration of Helsinki. The NIHR BioResource projects were approved by Research Ethics Committees in the UK and appropriate national ethics authorities in non-UK enrolment centres.

## Abbreviations

BCR: (B-cell receptor)
CADD: (combined annotation dependent depletion)
CID-G/A: (combined immun-odeficiency with granuloma and/or autoimmunity)
GWAS: (genome-wide association studies)
HGMD: (human gene mutation database)
*M*_*r*_: (mutation rate)
MRF: (mutation rate residue frequency)
PID: (primary immunodeficiency)
pLI: (probability of being loss-of-function intolerant)
*RAG1*: (recombination activating gene 1)
*R*_*f*_: (residue frequency)
RNH: (RNase H)
RSS: (recombination signal sequence)
SCID: (severe combined immunodeficiency)
TCR: (T-cell receptor)

## Authorship Contributions

Dylan Lawless analysed data and conceived and wrote the manuscript, Hana Allen Lango analysed NIHR genomic data, James Thaventhiran provided genomic data, NIHR BioResource–Rare Diseases Consortium provided genomic data, Rashida Anwar wrote the manuscript, Jolan E. Walter provided clinical genomics data and wrote the manuscript, Jacques Fellay wrote the manuscript, Sinisa Savic conceived and wrote the manuscript.

## Supplemental

**Figure E1:**
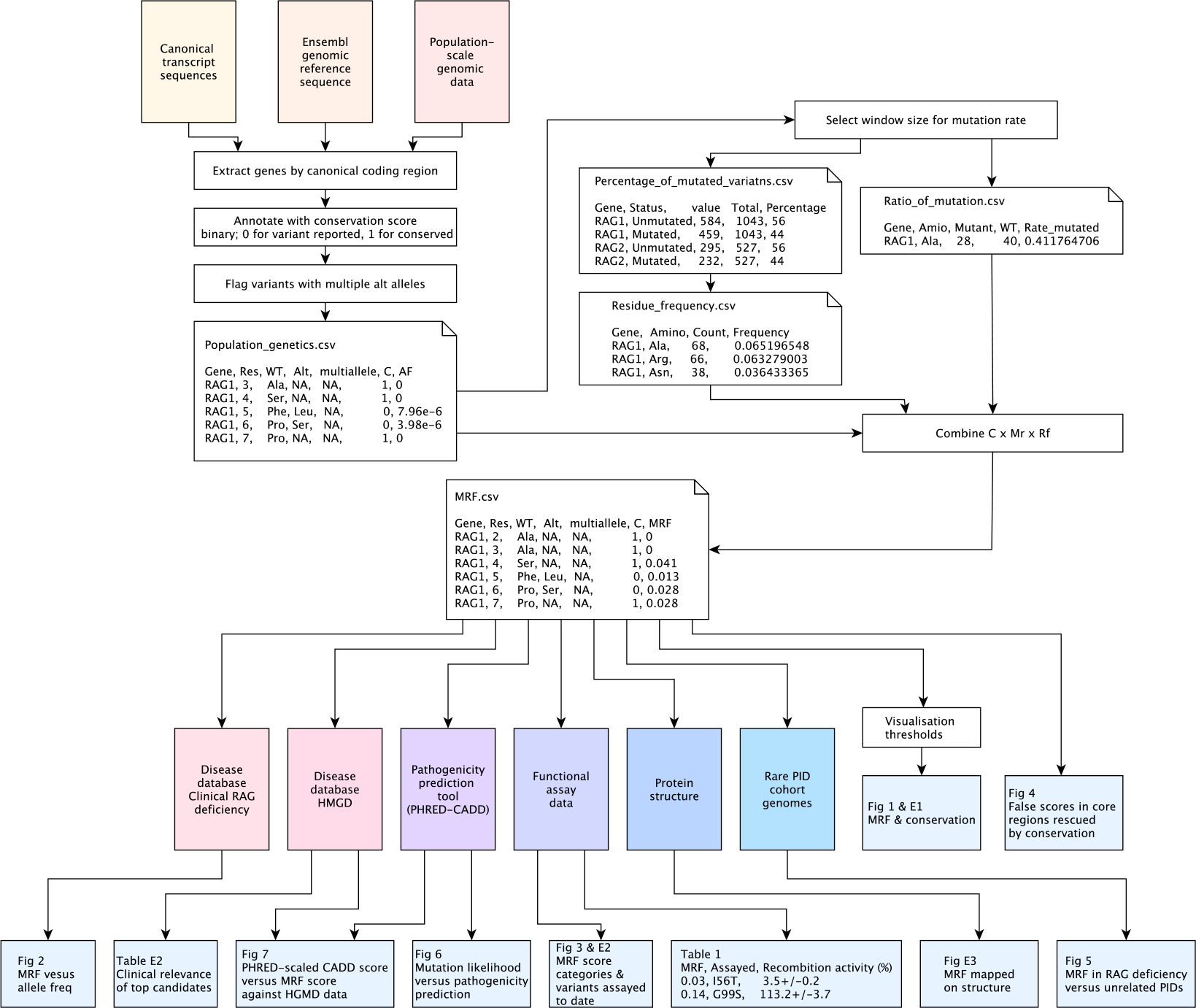
Data analysis summary map. Raw data and analysis scripts are provided in the supplemental. Analysis steps and data sources for each procedure described in *methods*. MRF; mutation rate residue frequency, PID; primary immunodeficiency.

**Figure E2:**
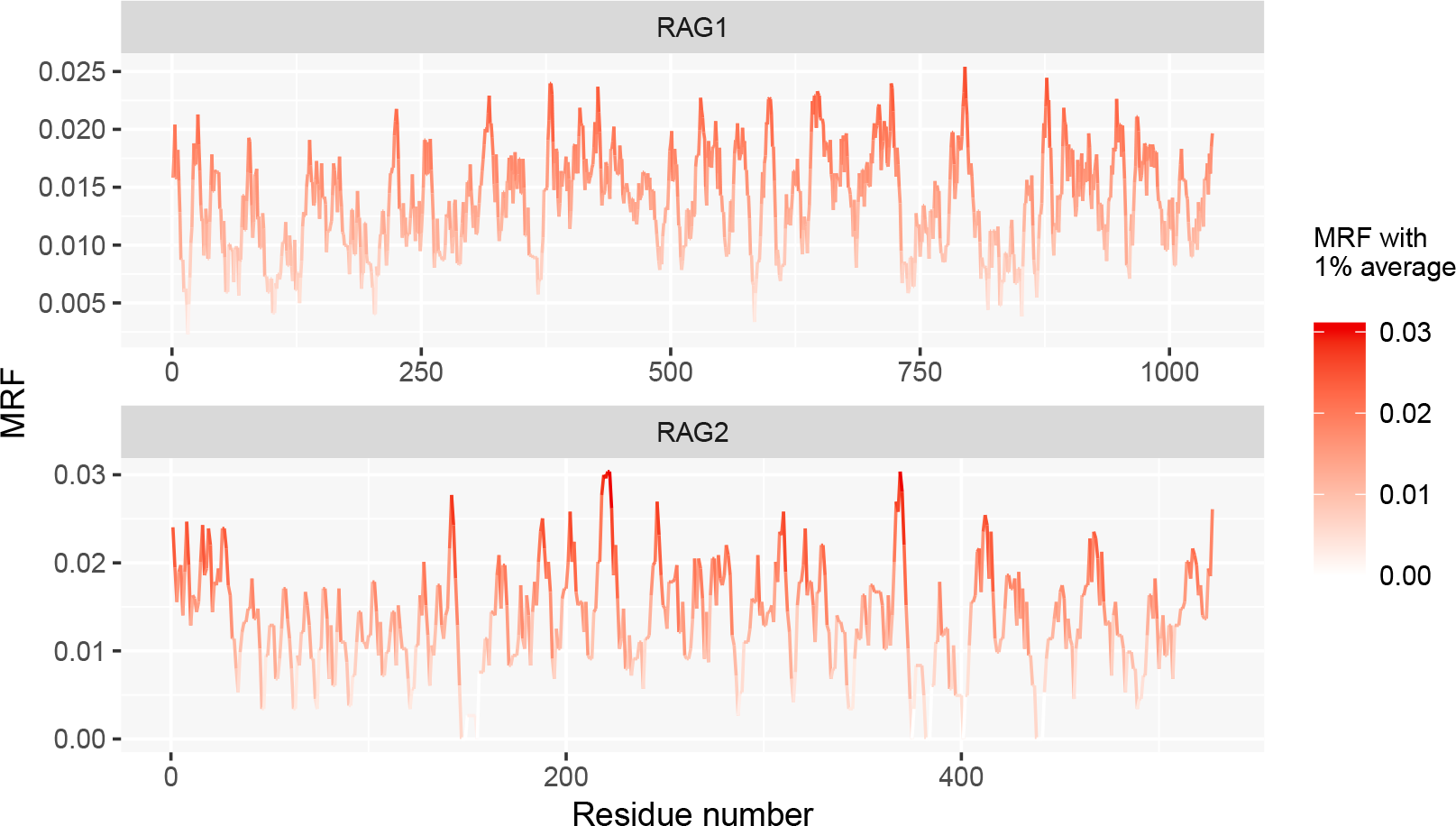
An alternative visualisation of MRF scores for RAG1 and RAG2 proteins. The data from Table E1 in column “Average over 1%” is displayed on both the y-axis and colour scale. An analysis-friendly long form CSV of the Table E1 data is also provided in the compressed supplemental R data “mrf.csv”.

**Figure E3:**
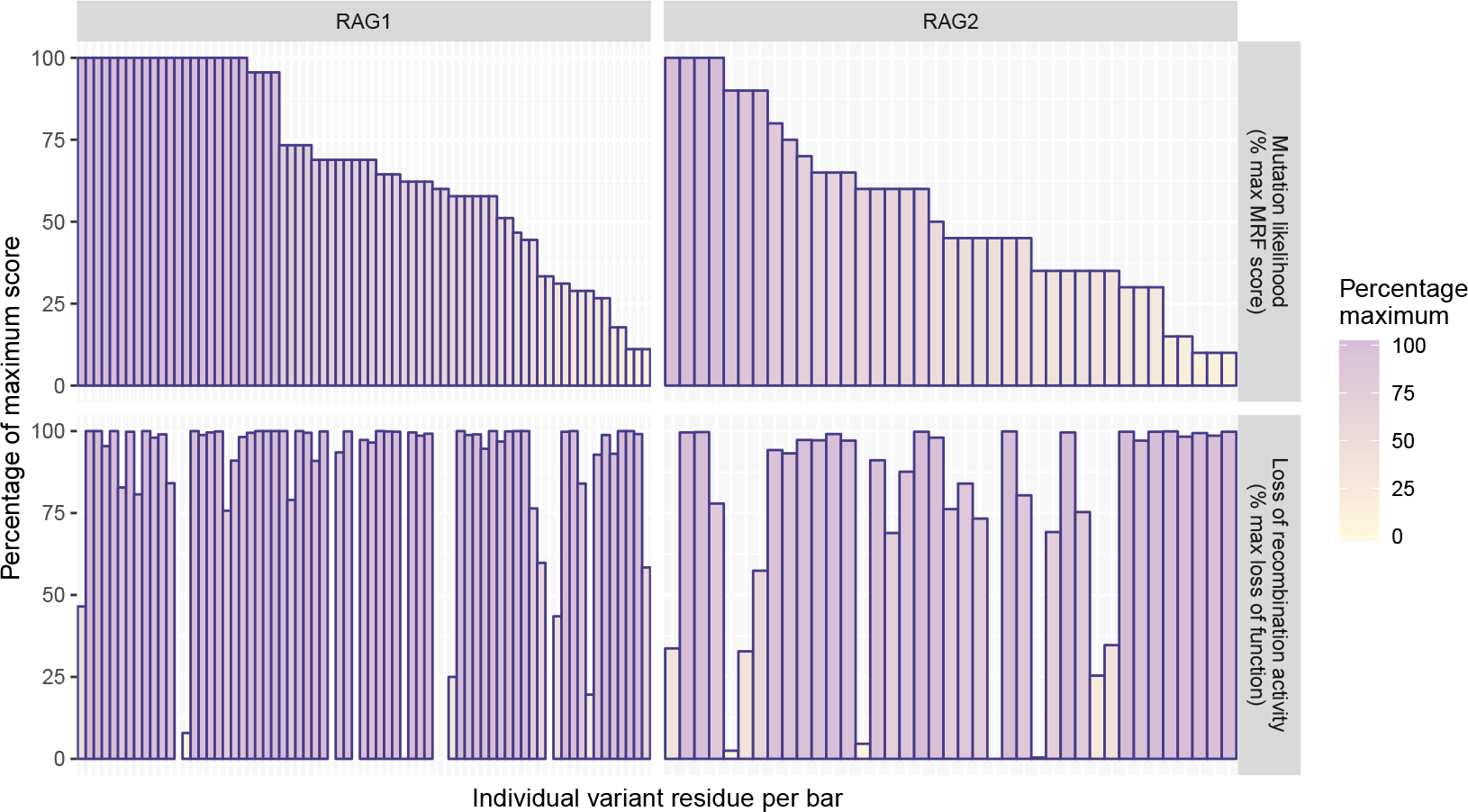
MRF likelihood score versus known functional activity. We compiled all variants that we know to have been assayed for protein function to date. The inverse of functional assay measurements were used, where 0% activity represents 100% loss of activity. MRF scores are presented as a percentage of the maximum score per gene (i.e., for RAG1 *MRF*_*max*_ = 0.043 (100%) and *MRF*_*min*_ = 0.0048 (0%)). **Top panels** show how likely each mutation is predicted to occur in humans. **Bottom panels** show the loss of protein activity as a percentage compared to wild-type (% SEM); most mutations tested produced severe loss of protein function, regardless of their mutation likelihood. Subset of *Recombination activity* data from Figure 3.

### Main supplemental data table

**Table E1:** MRF data tables. The complete scores are listed for each protein. Both the wild type and alternative variants reported on GnomAD are shown [21]. Multiple alternative variants are reported for some residues which are annotated in a separate column. A column listing the 1% average of MRF scores using a sliding window is present; this is used in Figure 1 (iii) with a cutoff threshold at the 75th percentile to clearly visualise high scoring clusters. The Boolean conservation score is based on population genetics data, where 1 represents no known variant at a residue site. The percentile values for 0.9 and 0.75 are provided to allow for alternative visualisation methods using the R source supplemental *“mrf.csv”* file.

### Clinical relevance of top candidates

The top scoring candidates in RAG1 were assessed for potential clinical relevance (Table E2). HGMD was chosen as a reliable, curated source of identifying pathogenic variants. 45% of RAG1 variants reported on HGMD (23 of 51) were predicted by our model as the most likely candidates seen clinically (the top scoring MRF group of had 66 residues total). The remaining variants in the top MRF group, which were not reported by HGMD (43 of 66), were assessed manually for their likelihood as potentially disease causing. 21 (49%) were highly conserved, not reported on GnomAD, and would be considered probable RAG deficiency on presentation as homozygous or compound heterozygous with a second damaging variant. The remainder had allele frequencies <0.0006, were only found as low frequency heterozygous in the general population, and justify functional validation. We expect that none of the top candidate mutations are benign.

**Table E2:** Clinical relevance of top candidates. 23 top MRF score variants were reported as pathogenic on HGMD to date. The remaining variants (the 43 not reported) were assessed by their frequency in population based on GnomAD (allele frequencies vary between individual variants but equate to approximately <6* and 9-77** heterozygous per 125,000 individuals). Therefore, no top candidates should be considered benign without functional validation. HGMD; Human Gene Mutation Database, MRF; mutation rate residue frequency.

| Most likely mutation candidates |  |
| --- | --- |
| Variant type | Number variants |
| Top MRF score candidates total | 66 |
| (i) Of which are reported on HGMD | 23 |
| (ii) Not reported on HGMD to date | 43 |

Table E2: Clinical relevance of top candidates.
| Top MRF score candidates not reported on HGMD |  |  |  |
| --- | --- | --- | --- |
| Frequency in Population | Number variants | Unreported top candidate (%) | GnomAD allele frequency |
| Not found | 21 of 43 | 49% | 0 |
| Very rare | 15 of 43 | 35% | <0.00002* |
| Very rare | 7 of 43 | 16% | <0.00006** |

### Supplemental analysis tables

**Table E3:** Known damaging in human cases. Amino acids and residue numbers are listed along with their basic mutation rate residue frequency value for known RAG deficiency [54]. Supplemental CSV file.

**Table E4:** Percentage of variants per gene. Percentage of mutated versus non-mutated amino acids in RAG1 and RAG2 based on GnomAD population genetics data [21]. Supplemental CSV file.

**Table E5:** Residue frequency. Basic statistics for RAG1/2 were produced using SMS2 [24]. Results are shown for the canonical sequences, a 1043-residue sequence of RAG1-201 peptide ENSP00000299440 and for a 527-residue sequence of RAG2-201 peptide ENSP00000308620. Both percentage and frequency of residue usage are provided. Supplemental CSV file.

**Table E6:** Simplified residue frequency. The data from Table E5 is simplified for use in data analysis to only include residues, count, total, and frequency per protein expressed. Supplemental CSV file.

**Table E7:** Ratio of mutation per gene. The number of times each residue is mutated was found in population genetic data. The number of mutant versus wild-type is shown, from which the rate of each is derived. Supplemental CSV file.

**Table E8:** Basic MRF scores. The mutation rate and frequency is shown which were used to calculate the basic mutation rate residue frequency for the main analysis dataframe and Table E1. Supplemental CSV file.

### Protein structure application

With the availability of a structured protein complex, modelling can be carried out prior to functional assays. Residues with the highest MRF for both RAG1 and RAG2 were mapped in Figure E4.

**Figure E4:**
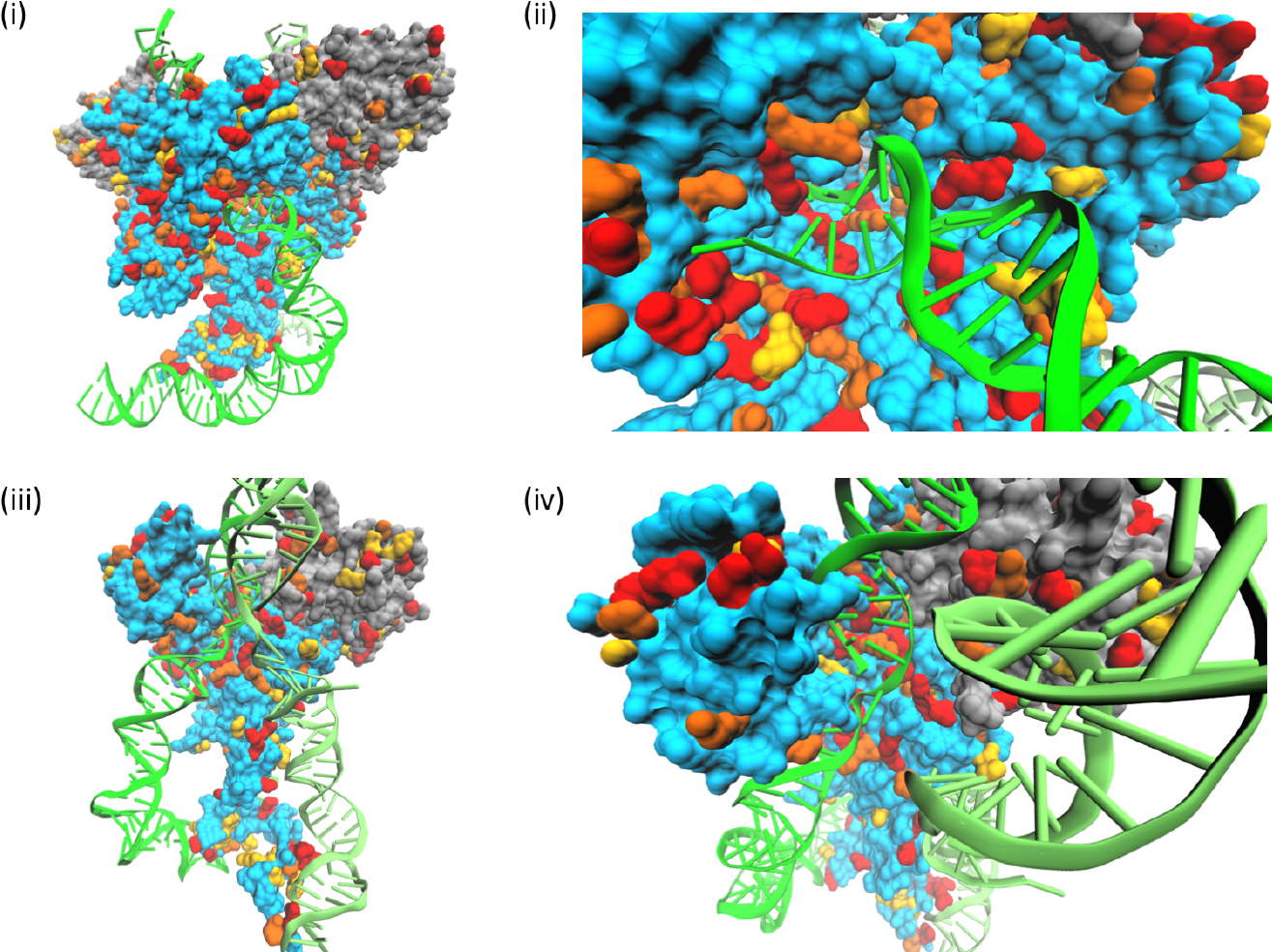
The RAG1 (blue) and RAG2 (grey) protein structure with top candidate MRF scores. (i) Protein dimers and (ii=iv) monomers illustrating the three highest category MRF scores for predicted clinically relevant variants. Increasing in score the top three MRF categories (illustrated in Figure 3) for each protein are highlighted; yellow, orange, red. DNA (green) is bound by the RAG protein complex at recombination signal sequences (PDB:3jbw).

### Median CADD score per residue

The sourced PHRED-scaled CADD score data consisted of nucleotide level values. We were interested in CADD scores averaged per codon. For every nucleotide position there were three alternative variants to consider, e.g.

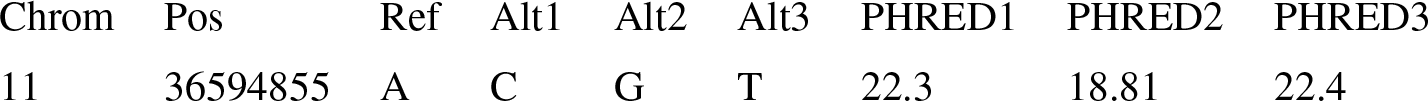

The PHRED-scaled scores are listed here; raw CADD scores are also included in the original database. To produce a working input we used the median score per codon, that is three scores per nucleotide and three nucleotides per codon. This produced median PHRED-scaled score per codon / residue, e.g.:

| Chrom | Pos | PHRED1 | PHRED2 | PHRED3 |
| --- | --- | --- | --- | --- |
| 11 | 36594855 | 22.3 | 18.81 | 22.4 |
| 11 | 36594856 | 25.3 | 23.6 | 24.6 |
| 11 | 36594857 | 24.8 | 24.3 | 24.5 |

Median PHRED = 24.3

#### Supplemental file

*‘RAG1.cadd.amino.csv’* within the analysis data *‘Raw_data_R_analysis_for_figures’* contains the median values over a three-nucleotide window, starting at nucleotide 1 to produce input data with the correct reading frame. The “PHRED-scaled” values are used as a normalised and externally comparable unit of analysis, rather than raw CADD scores. The area under the curve was calculated for density plots to quantify the difference between pathogenic and unreported variants with high scores, above the intersects >0.0409 and >22.84 for MRF and CADD, respectively, using score value (*x*) versus density (*y*) (Fig. 7 (i-ii)) with 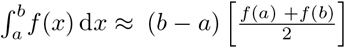.

### Genome-wide and disease-specific application

Weighting data can also be applied to the MRF score model to amplify the selectivity. The mutation rate can be applied genome wide with a process common in the study of information retrieval; term frequency, inverse document frequency (*tf* − *idf*). In this case the “term” and “document” are replaced by amino acid residue *r* and gene *g*, respectively such that,

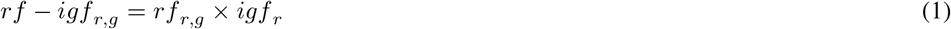

We may view each gene as a vector with one component corresponding to each residue mutation in the gene, together with a weight for each component that is given by (1). Therefore, we can find the overlap score measure with the *rf* − *igf* weight of each term in *g*, for a query *q*;

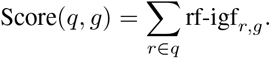

In respect to MRF scoring, this information retrieval method might be applied as follows; the *rf* − *igf* weight of a term is the product of its *rf* weight and its *igf* weight 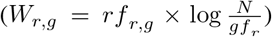 or 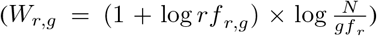. That is, firstly, the number of times a residue mutates in a gene (*rf* = *rf*_*r,g*_) and secondly, the rarity of the mutation genome-wide in *N* number of genes (*igf* = *N/gf*_*r*_). Finally, ranking the score of genes for a mutation query *q* by;

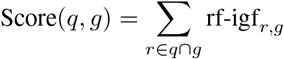

The score of the query (Score(*q, g*)) equals the mutations (terms) that appear in both the query and the gene (*r* ∈ *q* ∩ *g*). Working out the *rf* − *igf* weight for each of those variants (*rf*.*igf*_*r,g*_) and then summing them (∑) to give the score for the specific gene with respect to the query.

### Bayesian probability

MRF score may provide a limiting component required for applying Bayesian probability to disease prediction. A clinician may ask for the likelihood of RAG deficiency (or any Mendelian disease of interest) for a patient given a set of gene variants *P*(*H*|*E*) using Bayes’ theorem,

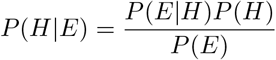

where *P*(*H*) is the probability of a patient having RAG deficiency, *P*(*E*|*H*) is the probability of RAG deficiency due to a set of variants that have been pre-emptively assayed, and *P*(*E*) is the probability of having a set of gene variants.

*P*(*H*) is known since the rate of RAG deficiency is estimated at an incidence of 1:181,000 [68], SCID at a rate of 1:330,000 [2], and we also recently show the rate of RAG deficiency in adults with PID [16]. Being a recessive disease, *P*(*E*) must account for biallelic variants and is the most difficult value to determine. This may be found from population genetics data for (i) the rate of two separate, compound heterozygous variants, (ii) the rate of a homozygous variant or potential consanguinity, or (iii) the rate of de novo variation [21]. *P*(*E*|*H*) would be identified where all variants are functionally validated. This requires a major investment, however the MRF score provides a good approximation.

